# Homology-based loop modelling yields more complete crystallographic protein structures

**DOI:** 10.1101/329219

**Authors:** Bart van Beusekom, Krista Joosten, Maarten L. Hekkelman, Robbie P. Joosten, Anastassis Perrakis

## Abstract

Inherent protein flexibility, poor or low-resolution diffraction data, or poor electron density maps, often inhibit building complete structural models during X-ray structure determination. However, advances in crystallographic refinement and model building nowadays often allow to complete previously missing parts. Here, we present algorithms that identify regions missing in a certain model but present in homologous structures in the Protein Data Bank (PDB), and “graft” these regions of interest. These new regions are refined and validated in a fully automated procedure. Including these developments in our PDB-REDO pipeline, allowed to build 24,962 missing loops in the PDB. The models and the automated procedures are publically available through the PDB-REDO databank and web server (https://pdb-redo.eu). More complete protein structure models enable a higher quality public archive, but also a better understanding of protein function, better comparison between homologous structures, and more complete data mining in structural bioinformatics projects.

**Synopsis:** Thousands of missing regions in existing protein structure models are completed using new methods based on homology.

## Introduction

Protein structure models give direct and detailed insights into biochemistry (Lamb *et al.*, 2015) and are therefore highly relevant to many areas of biology and biotechnology (Terwilliger & Bricogne, 2014). For decades, crystallography has been the leading technique to determine protein structure models (Berman *et al.*, 2014) and by now, over 120,000 crystallographic structure models are available from the Protein Data Bank (PDB) (Burley *et al.*, 2017). It is important to realize that all structures are interpretations of the underlying experimental data (Lamb *et al.*, 2015; Wlodawer *et al.*, 2013) and that the quality of a structure model should therefore be scrutinized by validation (Read *et al.*, 2011; Richardson *et al.*, 2013).

Due to numerous improvements in refinement and validation methods, the quality of protein structure models is continuously increasing (Read *et al.*, 2011): however, the completeness of models is decreasing (Fig. 1). About 70% of all crystallographic protein structures have missing regions (Djinovic-Carugo & Carugo, 2015) and this percentage is increasing. Typically, but not necessarily, these missing regions are loops between helices and strands. Loops occupy a large conformational space and therefore can be missing due to intrinsic disorder. Then, they cannot be modeled reliably in a single conformation. However, there are many cases where the experimental data give useful information on the loop conformation and hence many loops can be built into protein structure models. The term “loop” will be used in this paper in its broader definition to denote a missing region of protein structure, regardless of secondary structure conformation.

**Figure 1:**
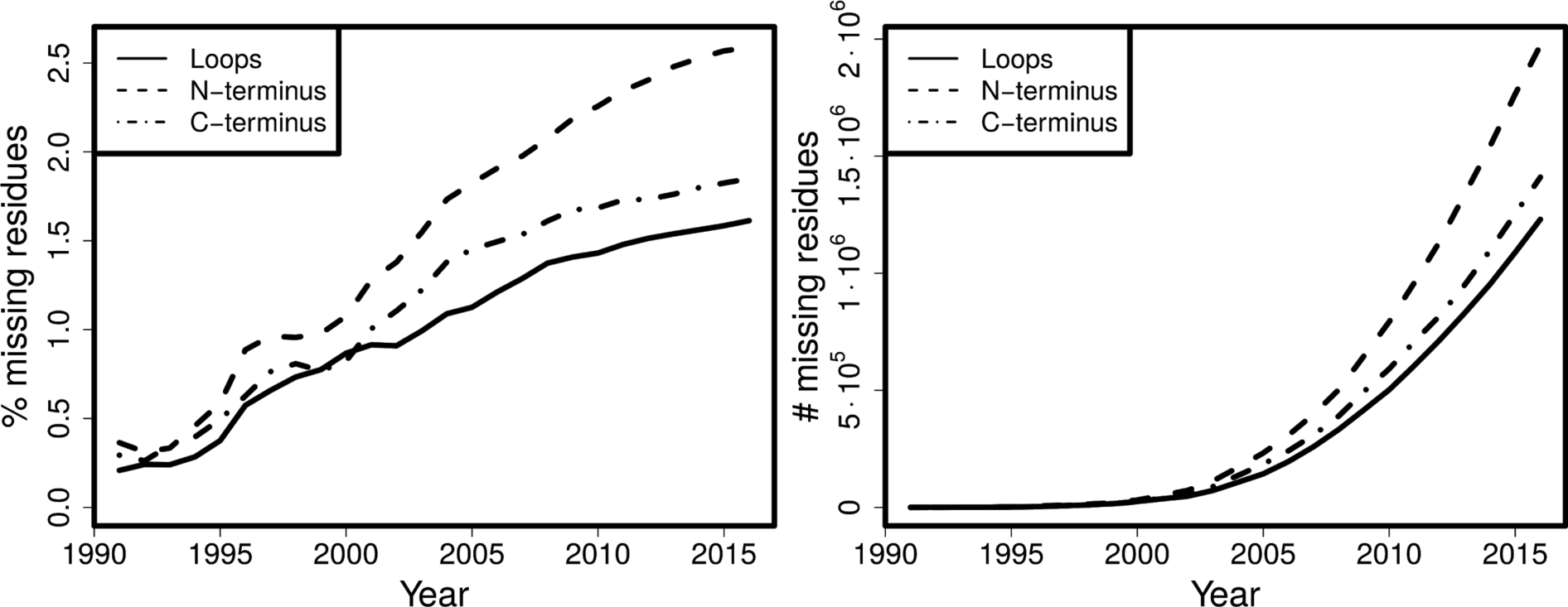
Cumulative percentage (left) and absolute number (right) of residues missing in termini and in loops for all structures over the years.

Missing protein regions or “loops”, are typically modelled towards the end of crystal structure determination. By that stage, all obvious features have typically been modelled, the electron density maps have improved, and it is possible to model the loops. Many programs are available to model loops either interactively (Emsley *et al.*, 2010) or automatically (Terwilliger *et al.*, 2008; Joosten *et al.*, 2008; Cowtan, 2012; Kleywegt & Jones, 1998; Depristo *et al.*, 2005). Completing the protein structure by modeling all loops that can be modeled, has two advantages: locally, the density becomes unavailable for modeling erroneous structural features such as other parts of the protein (e.g. side chains), crystallization agents, and water molecules; and globally, a correctly fitted loop will reduce the phase error and give an overall improvement of the electron density maps. Available loop-building approaches rely on one form or another of conformational sampling that attempts to find the best fitting conformation of the loops based on the local electron density and our general knowledge of protein structure.

Building loops is often one of the most difficult, time-demanding, and sometimes frustrating stages of crystallographic model building. If loops are too disordered to yield traceable electron density, they cannot be built and there is no problem. In many cases however, loops are sufficiently rigid to yield interpretable density. Typically, this electron density is not as clear as desired, which makes it challenging for crystallographers and model building programs to model a loop in a realistic conformation. Whether this is eventually successful depends on many factors, such as perseverance and skill of the crystallographer(s) and the algorithmic quality and ease-of-use of loop building programs. Noteworthy are algorithms that attempt to interpret the electron density with multiple loop conformations (Burnley *et al.*, 2012; Levin *et al.*, 2007). While there is little doubt that very often multiple conformations can represent the experimental data better, here we wanted to deal with the problem that a loop is not modelled at all despite strong experimental evidence that it can be modeled.

Previously, we developed algorithms to transfer information about homologous protein structures for obtaining geometric restraints for low resolution refinement (van Beusekom *et al.*, 2018). Here, we exploit the relation between homologous protein structures for loop building. We reasoned that as highly similar protein structures have very similar structure, the conformation of a loop in one homologous structure can assist in identifying the loop location in another homologous structure, that is being built. The presence of a loop in a homologous structure is also an indication that loop building is viable. If a region has not been modelled in any of the homologs, it is unlikely to be buildable in the new structure (provided there are many homologous models solved by different crystallographers), but if it is built at least once there is a good chance that the same region can be modeled in the structure in question. Of course, there are exceptions to the similarity of homologous loops, as loops can adopt distinct conformations which are often of high significance for function: there are countless examples that describe loops motions that are associated with interaction with a ligand or change in the context of different crystallographic contacts.

It has already been shown for a set of 16,370 PDB structures (Le Gall *et al.*, 2007) that while 92.2% of all crystallized residues was always ordered in all homologs and 4.4% was always disordered, 3.3% is “ambiguous”: these residues were modeled in at least one structure but disordered in other(s). Another survey (Zhang *et al.*, 2007) observed that such regions, named “dual personality” fragments, occur in 45% of sequence-identical structure groups, and within those groups, they occur almost twice on average. Of the ambiguous loop regions, 59% are predicted to be ordered by all three protein disorder predictors used in a third study (DeForte & Uversky, 2016). Thus, the increasing redundancy of homologs in the PDB, the increasing percentage of unmodeled residues (Fig. 1), and the disorder predictions for these regions, argue that using “dual personality” regions between homologous structures, is a viable strategy for increasing the completeness of protein structure models.

We have therefore decided to develop a procedure for building loops in cases that a loop is available in other homologous structure(s). The implementation of these new ideas, has taken place in the context of PDB-REDO, a procedure we are developing to (re-)refine and partially rebuild protein structure models, both retrospectively (by updating existing PDB entries (Joosten *et al.*, 2012)) and proactively (our software is available as a web-server (Joosten *et al.*, 2014) and for local installation). As the PDB-REDO electron density maps are often better than the original maps that can be simply recalculated from the PDB entries, incorporating this work in the PDB-REDO framework allows to have the best possible maps for the building and validation of the missing loops. For every structure in question, the missing loop is first identified, then built by grafting it from a homologous structure, refined to fit the electron density map in real space, and finally validated against geometrical criteria and the electron density. We have built and validated several thousands of loops missing in structures deposited in the PDB. Here we discuss the methods and show some examples where our procedure makes a notable difference in the structure model.

## Methods

### Loop building

We have developed algorithms to transfer loops from homologous protein structures to the target structure (Fig. 2). The manner of handling the homologs in *Loopwhole* is nearly identical to our previous program *HODER (van Beusekom et al., 2018)*, which generates homology-based H-bond restraints. The only difference is that homologs are not filtered by resolution in *Loopwhole*. The default maximum length for attempted loop transfer is 30 amino acids.

**Figure 2:**
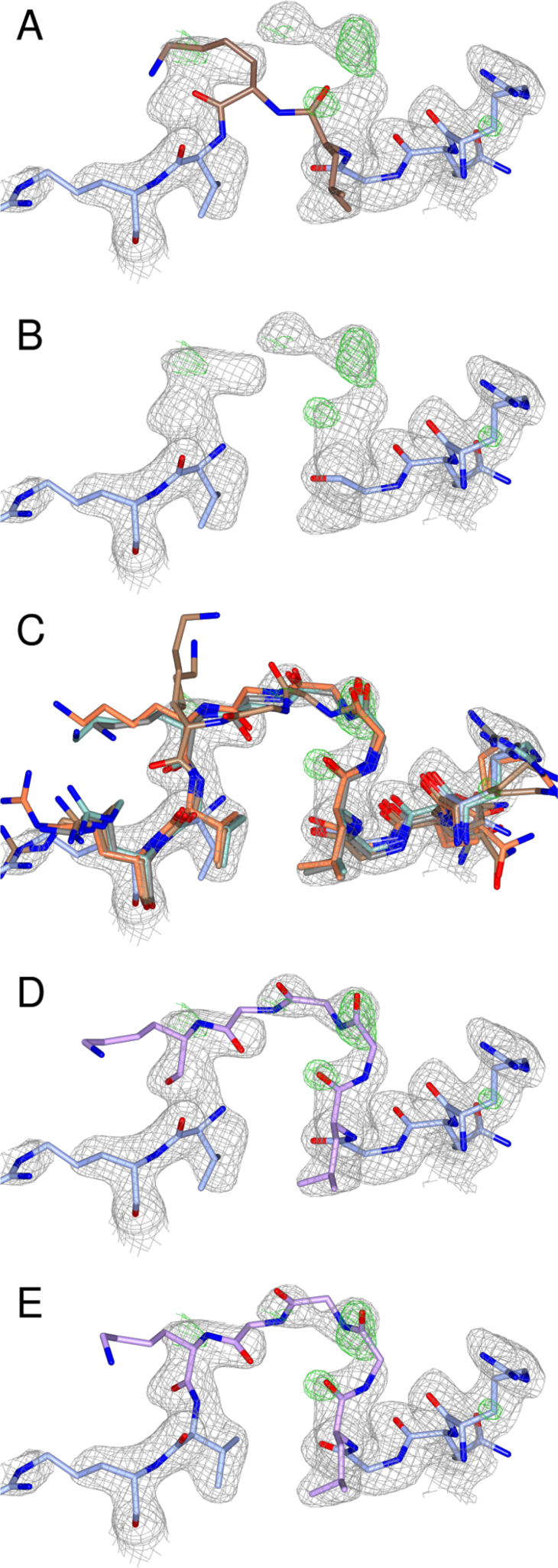
A stepwise illustration of the Loopwhole algorithm, applied to three missing glycines (PDB entry 1dmn (Kim et al., 2000)). All 2mF_o_ - DF_c_ and mF_o_ - DF_c_ electron density maps are shown at 0.7 σ and 3.0 σ, respectively. (A) The PDB structure has no apparent gap: two residues are wrongly bound to one another (highlighted in brown). The gap is detected from sequence. (B) The two residues immediately adjacent to the missing loop are removed, as these are often in the wrong conformation. (C) All homologous chains with the loop present are structurally aligned to the target structure model (only four homologs shown). (D) If the surrounding residues align well, the loop is grafted into the target structure model. By default, the top ten alignments are kept (only one shown). (E) After real-space optimization, the loop with the best density fit is kept provided the density fit and geometrical quality pass the filter criteria.

Before any loop building is attempted, the density fit is evaluated per chain with EDSTATS (Tickle, 2012). In the rare case that the RSCC for an entire chain is below 0.80, we do not attempt to build loops in that chain, but instead warn the user that the density fit is low which makes the overall chain conformation unreliable.

There are two initial requirements for loop transfer: the presence of unmodeled loops in the input structure and modelled equivalent loops in the homologs. Unmodeled loops are detected using PDB-REDO’s *pdb2fasta*, and high-identity homologs are aligned by sequence. The details of these algorithms have been described in earlier (van Beusekom *et al.*, 2018). If both requirements are met, loop transfer is attempted for each of the homologs that has a complete backbone model for the homolog. Both input model and homolog are required to have at least five consecutive modeled residues on each side of the loop. Of these ten residues, the two residues directly adjacent to the loop are remodeled together with the loop because they were often found to be in a suboptimal conformation. The remaining eight residues are used for alignment. To prepare residues for the alignment, side-chain atoms are deleted in case of mutations and administrative flips for Asp, Glu, Phe and Tyr residues (DEFY flips) are done to ensure equivalent atoms are in equivalent positions. A homolog is skipped if the sequence identity in the loop or in the aligned residues is less than 50%; the exception is a single-residue loop, which is allowed to be mutated. Finally, structural alignment is performed using quaternions (Kuipers, 2012).

If the backbone RMSD of alignment is less than 2.0 Å (in default settings), an initial loop transfer is performed. The two residues directly adjacent to the loop are deleted and the aligned loop, including the aligned two directly adjacent residues, is inserted into the protein model. In the transferred loop, side-chains are cropped where appropriate in case of mutations, the occupancy of all atoms is set to 1.00 and the B-factor is multiplied for each atom by the ratio of average B-factors between input structure and homolog. Methionine is mutated to selenomethionine and *vice versa* based on the other residues present in the input model.

The next check for the initial transfer of a loop is the evaluation of clashes with the modeled atoms already present, or with symmetry copies thereof. We make a distinction between main-chain and side-chain atom clashes. Since the position of a Cβ is significantly limited by the main-chain conformation, we include this side-chain atom in the main-chain for clash analysis. Heavy clashes are defined as atom-atom distances <2.1 Å and small clashes as <2.6 Å.

In case of clashes, important atom(s) must be retained. In *Loopwhole* we use hierarchical rules in atom importance to decide how to proceed. The atoms that are always kept are main-chain atoms and most ligands (the exceptions are glycerol, ethanol, and 1,2-ethanediol and its polymeric (PEG) forms). Whenever a main-chain atom from a loop candidate clashes with the previously-modeled backbone or most ligands, the loop candidate is discarded. In contrast, if the main chain of a loop candidate clashes with a compound from the list of exceptions (such as glycerol), that compound is discarded. The second most important group are the side-chains: they can be discarded temporarily to be added back later by the program *SideAide (Joosten et al., 2011)*. Previously-modeled side-chains are considered more important than loop side-chains. Side-chains, in turn, are considered more important than water molecules and any atoms with an occupancy of 0.01 or lower. These principles led to the following decisions.

Previously-modeled side-chains are removed only if they clash heavily with the loop backbone, unless they form a cysteine bridge: in such cases the loop candidate is discarded. Loop side-chains (from γ-atom(s) onwards) are deleted if they clash heavily with any previously modeled protein (main-chain or side-chain) or with any other compound except for water. Ligands from the exception list are removed whenever they clash heavily with the main chain of the loop. Waters and atoms with an occupancy of 0.01 or less are removed even in case of small clashes with any loop atom.

If there are no insuperable clashes, the loop candidate is saved and existing candidates are sorted by RMSD. If two loop candidates have a very similar conformation (RMSD < 0.1 Å), the candidate with the worst RMSD of alignment is discarded. Once all BLAST hits are evaluated, the top candidates (by default, the top ten) are subjected to real-space refinement by *coot-mini-rsr (Emsley et al., 2010)* using torsion angle restraints. One extra residue from the existing protein on either side of the loop is added to the real-space refinement region, which allows the existing protein model to better adapt to the new loop. In the *coot-mini-rsr* input PDB file, clashing atoms are removed, atom numbering is updated (including CONECT records) and ‘gap’ LINK records are deleted. Sometimes, there are still small gaps at the boundary of the transferred loop and the existing model. To increase the success rate of *coot-mini-rsr* of closing these gaps, the backbone nitrogen atoms on the loop edges are moved into this gap. This can temporarily create unlikely atom bond lengths and angles, but these will be resolved in real-space refinement (or in the subsequent reciprocal-space refinement).

After running *coot-mini-rsr*, it is checked again if there are no insuperable clashes between the loop and the protein, because we observed the loop may be placed into density of other moieties in real-space refinement, such as a symmetry copy of itself.

At this stage, all remaining loop candidates with bad geometry are discarded. First, candidates where there is no peptide bond between two consecutive residues or where *coot-mini-rsr* has not converged to a minimum are removed. The resulting RMS-Z scores from *coot-mini-rsr* are used to filter bad geometry candidates: loops are rejected if bond or angle RMS Z exceeds 1.2, chirality RMS Z exceeds 1.5, or if plane or torsion RMS Z values exceed 2.0. In this filter the RMS-Z values are allowed to be relatively high because subsequent reciprocal-space refinement will further improve the loop. Loop candidates are only allowed to have *cis*-peptides if the corresponding residue in the original loop in the homolog is also a *cis*-peptide. Loops that have multiple sequential distorted omega angles (maximum deviation 30° from 0° or 180°) are discarded, but single distortions are allowed as these are usually resolved in subsequent refinement. Finally, loop candidates are evaluated on their Ramachandran Z score. If the Z score is poor (lower than −5), it is compared to the Z scores of the other loop candidates and also to the Z scores of the loop in the homologous structure models from which it was adapted. Then, the candidate is discarded if it is a 2σ negative outlier (according to the Grubbs’ test (Grubbs, 1950)), either compared to the other loop candidates or compared to the original conformation of the loop. The Ramachandran Z-score calculation is performed using the algorithms of the new PDB-REDO program *tortoize*. This algorithm is based on the implementation in WHAT_CHECK (Hooft *et al.*, 1997) and is described in the Supporting Information.

The density for each remaining candidate is then computed using cubic interpolation function from clipper (Cowtan, 2003). It is computed only for the main-chain atoms of the candidate loop to ensure that the metric is not influenced by the presence, absence, or length of the side-chains of the loop. Additionally, the density is computed for all main-chain atoms that are ordered in all homologs, *i.e.* the set of atoms that are always ordered. If there are fewer than 30 atoms in this set, all non-loop main-chain atoms in the input structure model are taken. The ratio between average loop candidate density and the average density of the control set is then computed. This ratio must be over 0.25 for a loop candidate to be acceptable. The cut-off was established after manual inspection of several hundred candidate loops.

Finally, there is an option to subject a number of candidates (by default only the loop with the best density fit) to the PDB-REDO programs *SideAide* and *pepflip (Joosten et al., 2011)* to complete the side-chains and check for potential peptide flips of the loop area. However, this is not default behaviour since these programs are already run after *Loopwhole* in the PDB-REDO implementation. However, *Loopwhole* writes a list of amino acids whose side chain is incomplete: at low resolution, *SideAide* is not run by default on all amino acids in PDB-REDO, but only on amino acids in a list, to which the novel residues in the loop are added.

After the optional running of *SideAide* and *pepflip*, the loop with the best main-chain best density fit is kept.

There are a few special cases where detection or building of a missing loop is more complicated. First, there is the possibility that a loop is in fact modeled, but with all atoms modeled at zero occupancy, or at occupancy 0.01. We consider such loops as unmodeled and proceed as described above, however, we treat the current zero-occupancy loop itself as an extra candidate. Since this loop is already at the correct location in the model, no alignment is necessary; apart from that, this candidate is treated the same as others.

Another special case is dealing with alternates in and near loops. If there are any main-chain alternates among the residues that are to be aligned with homologs, the missing loop is skipped because the alignment target is ambiguous. An exception is made if the only backbone alternate atoms are Cα atoms (which is common for residues with alternate side-chain conformations): then simply the first atom is picked. In such cases, the positions of alternate Cα atoms are very close to each other. Alternate side-chains are truncated before alignment. In homologs, backbone alternate conformations are treated as separate candidates: structure alignment is performed for each alternate in the homolog and/or each alternate loop is transplanted. If there are multiple stretches that contain alternates with full occupancy atoms in between, combinations of these alternates must be aligned for completeness and the exponential increase of combinations makes computation expensive, therefore these (rare) cases are excluded.

Finally, there are cases where residues next to a loop are only partly present. In such cases, the partly-modeled residue is also removed before the loop fitting. That is, the loop is extended by one more residue and the partial residue is replaced.

### Adding missing atoms, atom pairs, or atom trios: *fixDMC*

We observed that some protein models miss one or several atoms from a peptide backbone. In order to also correct these smaller missing parts, the program *fixDMC (fix “dat” main chain)* corrects these omissions, adds missing C-terminal oxygens, and resets occupancies to 1.0 in regions where there are no alternates and surrounding atoms are modeled at full occupancy.

We make use of the fact that C□^i^, C^i^, O^i^, N^i^, and C□^i+1^ lie in a plane. Whenever at least three atoms of a single plane are present, this planarity combined with the known geometry of an amino acid gives enough information to compute its coordinates. The C□ atom lies in two planes: that of the preceding and that of the following amino acid residue. Therefore, it can be added based on either of these residues. Applying the geometrical rules of the planarity extensively, we can compute any set of 1, 2, or 3 atoms provided the preceding and following residues are modeled.

Additionally, *fixDMC* uses functionality from PDB-REDO’s *pdb2fasta (van Beusekom et al., 2018)* to add the second C-terminal oxygen (“OXT”) if the SEQRES records or user-inputted FASTA file indicate that the complete C-terminal residue has been modeled except for this atom. The addition of this atom can also be based on the peptide plane.

Finally, the occupancy of protein atoms is reset to full occupancy if the residue contains no alternates and the preceding and subsequent atom are both modeled at full occupancy. An exception is made for the carbonyl oxygen: since this atom is only bound to a single carbon atom, only that carbon atom is required to be modeled at full occupancy.

### Implementation in PDB-REDO

The program *fixDMC* is run at the early stages of PDB-REDO after the initial electron density maps are calculated, before any individual atomic coordinate or B-factor refinement. The OXT atoms are only added if it is reliable that the final modeled residue is the actual C-terminus of the crystallized construct. Therefore this step is only performed if the header of the input PDB file has SEQRES records or if user-supplied sequence(s) can be mapped to the modelled atomic coordinates.

*Loopwhole* is run after the initial refinement in PDB-REDO. The default behaviour is to always attempt to build loops, but this can be switched off if needed. It should be noted that on the PDB-REDO web server loops can only be built if the sequence of the missing residues is known. That is, users must supply the sequence as a FASTA file or as SEQRES records in the PDB file. If *Loopwhole* builds any residues the electron density map is recalculated by *Refmac5 (Murshudov et al., 2011)* before the other rebuilding stages of PDB-REDO (Joosten *et al.*, 2011).

Sequence files in PDB-REDO mark residues with a complete backbone in uppercase letters and incomplete or unmodeled residues with lowercase letters (van Beusekom *et al.*, 2018)), an idea adopted from the SEQATOMS server (Brandt *et al.*, 2008). Therefore, both *Loopwhole* and *fixDMC* write updated FASTA files to reflect changes in residue completeness or presence. Additionally, *Loopwhole* updates the TLS groups in PDB-REDO. If a TLS group is surrounding the loop, the loop is added in that group; if the loop is on the border of two TLS groups, it is added to the first one.

At the final stage of PDB-REDO the program *Modelcompare* writes a data file that is used by COOT (Emsley *et al.*, 2010) and 3Dbionotes (Segura *et al.*, 2017; Tabas-Madrid *et al.*, 2016) to highlight the new loops.

### Testing

*Loopwhole* was run over all entries available in PDB-REDO to identify which loops could be built. Hundreds of randomly selected loops were manually analyzed to empirically establish the validation cut-offs mentioned above. Finally, from all entries in which *Loopwhole* built loops, 2,000 entries were randomly selected for further analysis in PDB-REDO. These entries were subjected to the PDB-REDO pipeline twice: once with and once without loop building. Due to various limitations (not related to loop building), 10 PDB-REDO jobs were not completed, hence the final test set consisted of 1,990 entries.

## Results

### Loop building

The computer program *Loopwhole* was developed to build protein loops based on homology (Fig. 2). We first applied *Loopwhole* on the structures available in the PDB (Table 1). When *Loopwhole* was then applied on the PDB-REDO databank, we observed an increase of 11% in the number of built loops. This is likely because the structure models and the electron density maps in PDB-REDO (which are obtained after modern re-refinement and re-building) are of higher quality than their “static” PDB counterparts. The total number of missing loops in the PDB-REDO data bank was 148,919. An initial loop was constructed by *Loopwhole* in 66,035 cases (44%). For the other 56%, there were either no homologous loops available, or the loop conformation was too different between the “donor” and “acceptor” structures (due to genuine structural differences, or because of “sequence register” errors. Another 41,073 loops (28% of total) were discarded according to various validation criteria (Fig. 3). Many loops were rejected as their fit to the electron density was too poor (Fig. 3B), and less often based on geometrical criteria or because both density fit and geometry were poor. The remaining loops have excellent geometry, typically better than the loop in the original structure (Fig. 3C) and a good fit to the density.

**Table 1:** Number of built loops and affected structures in the PDB and PDB-REDO databanks. We used 112,385 PDB-REDO entries available in February 2018.

| Model origin | # Loops built | # Entries modified |
| --- | --- | --- |
| PDB | 22,480 | 10,806 |
| PDB-REDO | 24,962 | 11,571 |

**Fig. 3:**
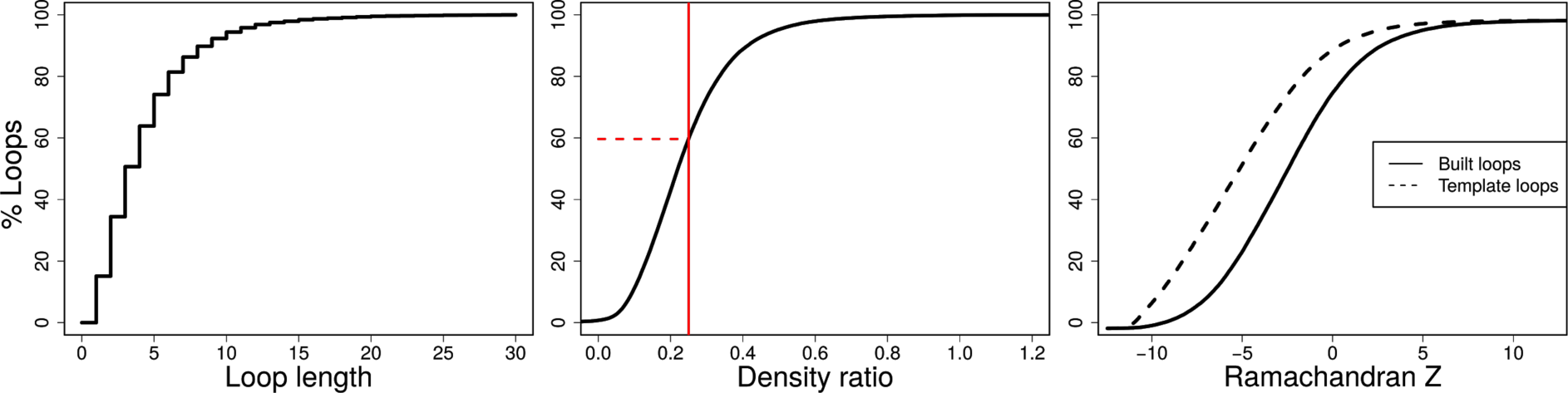
Cumulative distributions for properties of loops that can be built in the PDB. (Left) Most of the buildable loops are short. (Middle) The density ratio of loop candidates; in contrast to the other two subfigures this includes loops that were not built. The minimum required density ratio of 0.25 is indicated by the vertical red line. Of all initial loop candidates, 60% has insufficient density and is therefore discarded. (Right) The Ramachandran Z-score for candidate loops and their counterparts in the structure model from which they were taken. The backbone conformation of the built loops is excellent and better than the conformation of loops in structures from which they were taken, which is largely due to the application of Ramachandran restraints in the real-space refinement of loops in *coot-mini-rsr* (Emsley et al., 2010).

The current version of *Loopwhole* was able to build a total 24,962 missing loops in 11,571 entries. For 359 cases in which a loop was built, a zero-occupancy loop was present in the original model. To place the loops, 18,449 water molecules were removed; additionally, small molecules such as glycerol or ethanediol were removed in 22 cases. The distribution of the length of the built loops is shown in Fig. 3A.

Next, we wished to examine if incorporating *Loopwhole* in the PDB-REDO pipeline, impacted performance of PDB-REDO as a whole. We thus ran PDB-REDO on 1,990 randomly selected structures in which loops could be built, once with and once without loop building. The impact of loop building on standard validation metrics (Read *et al.*, 2011) such as the R_free_, Ramachandran Z score and packing Z score was minimal (Fig. S1), while the mean RSCC and RSR values (indicating the fit to the density) for the loops themselves were 0.75 and 0.14, respectively (Fig. 4). These values are, naturally, lower than for well-defined parts of the structure model, but are for example consistent with density criteria for acceptable ligands (Weichenberger *et al.*, 2013; Cereto-Massagué *et al.*, 2013; Warren *et al.*, 2012). From this we concluded that loop building in the context of PDB-REDO is useful, and this feature is already enabled by default. For practical reasons (lack of CPU time) the remaining PDB entries, are being gradually “redone” and placed in the PDB-REDO databank.

**Figure 4:**
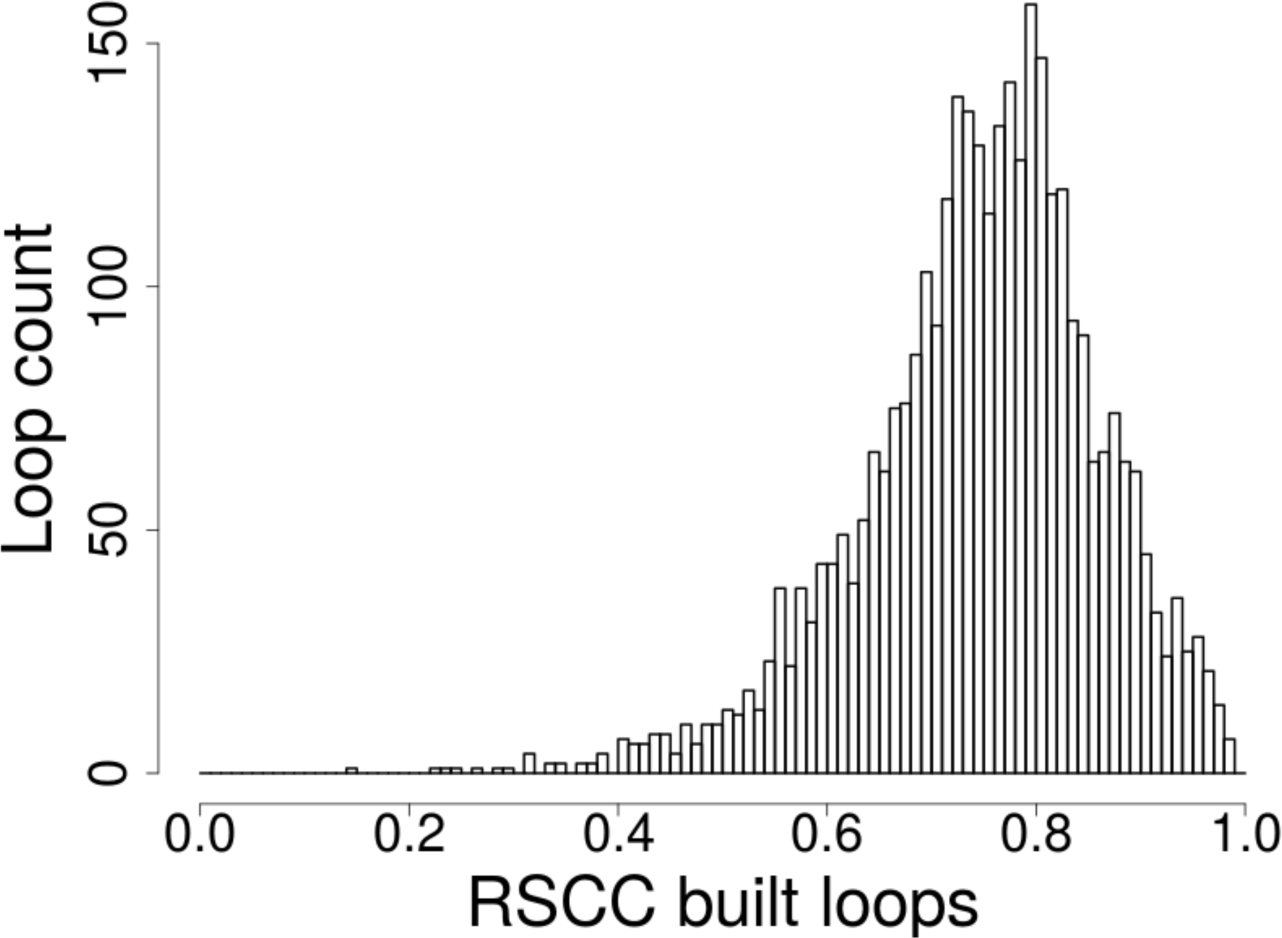
The distribution of the real-space correlation coefficient (RSCC) for all loops built in 1,990 PDB-REDO entries. The RSCC was calculated on the final PDB-REDO structure models.

### Completing the main chain

Rather surprisingly, we found that in many structures in the PDB there are missing individual atoms in the main chain. We therefore created the program *fixDMC* that can add one to three missing main chain atoms based on the geometry of existing atoms (see Methods for details). Running it in the same PDB-REDO dataset as above, single atoms were added in 1,500 cases (out of which 1,281 were carbonyl oxygens), atom pairs were added in 55 cases and in 40 residues, three atoms were added. Additionally, there were 38,101 cases with individual backbone atoms that had their occupancies set to values less than 1.00 without being part of an alternate conformation or of a partially occupied peptide ligand, out of which 2,926 cases had an occupancy of zero. Finally, we found that the second C-terminal oxygen atom (OXT in PDB nomenclature) is missing in many PDB entries; *fixDMC* added OXTs to 41,120 protein chains, were the terminal residue in the structure coincides with the terminal residue in the declared construct sequence. Notably, the percentage of protein chains with missing C-terminal oxygens has been steadily increasing over the years (Fig. 5). In 2017, in 44% of the chains with a modeled C-terminal amino acid, OXT is missing.

**Figure 5:**
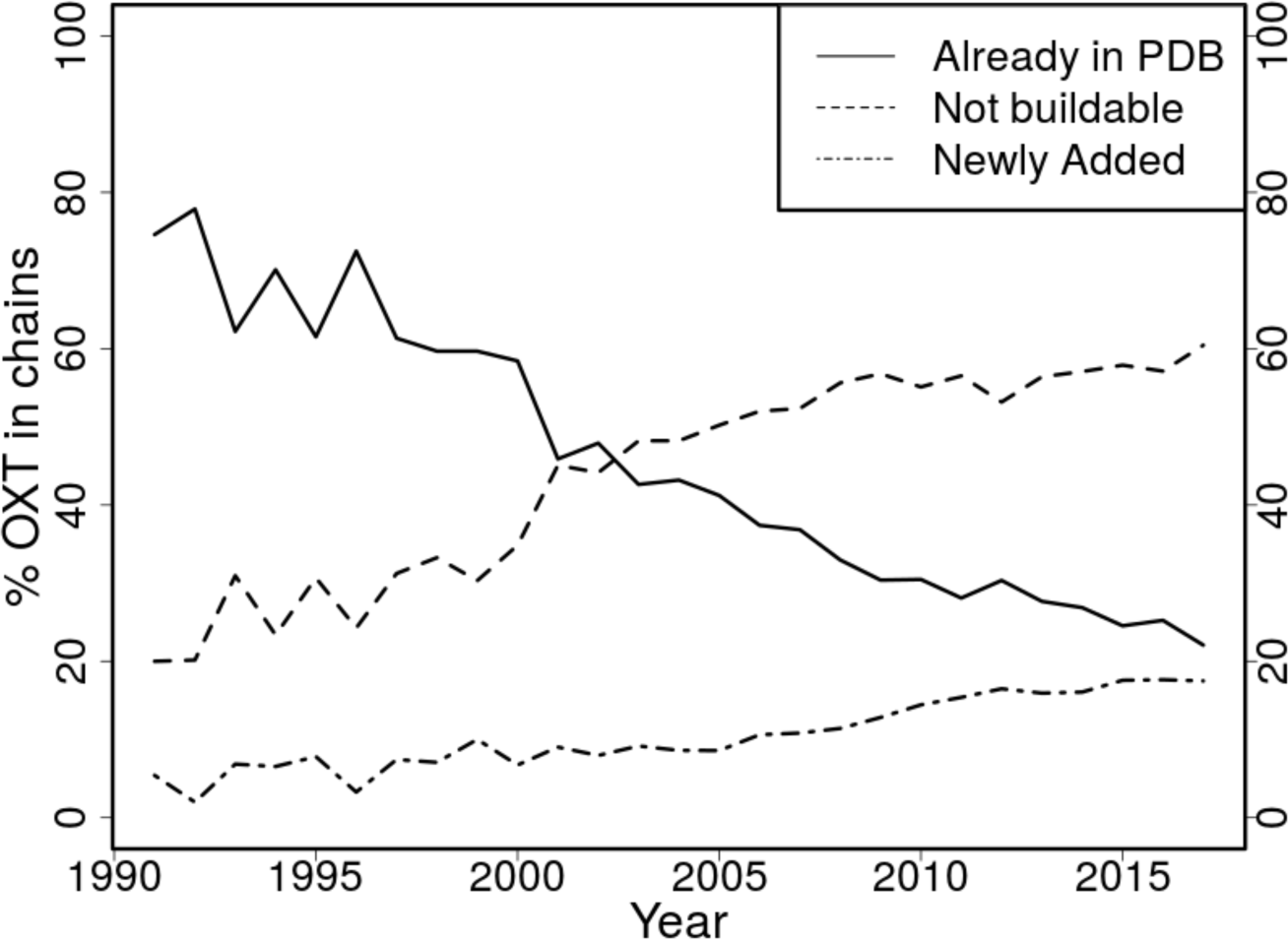
The percentage of C-terminal oxygens (OXT): present in the PDB; missing the C-terminal amino acid and thus not buildable; and newly added to the C-terminal amino acid.

### Examples of built loops

To illustrate the relevance of building loops in the PDB, we here show several examples in which *Loopwhole* clearly improves structure interpretation.

First, a structure of β-glucosidase (PDB entry 3abz (Yoshida *et al.*, 2010)) has seven missing regions, five of which can be added. Only the first of four NCS copies is complete. One of the missing regions, a stretch of 14 residues, is shown in Fig. 6A. The electron density for this loop is very clear. By adding the loops to the structure, the structure model is now much more complete and thus better interpretable.

**Fig. 6:**
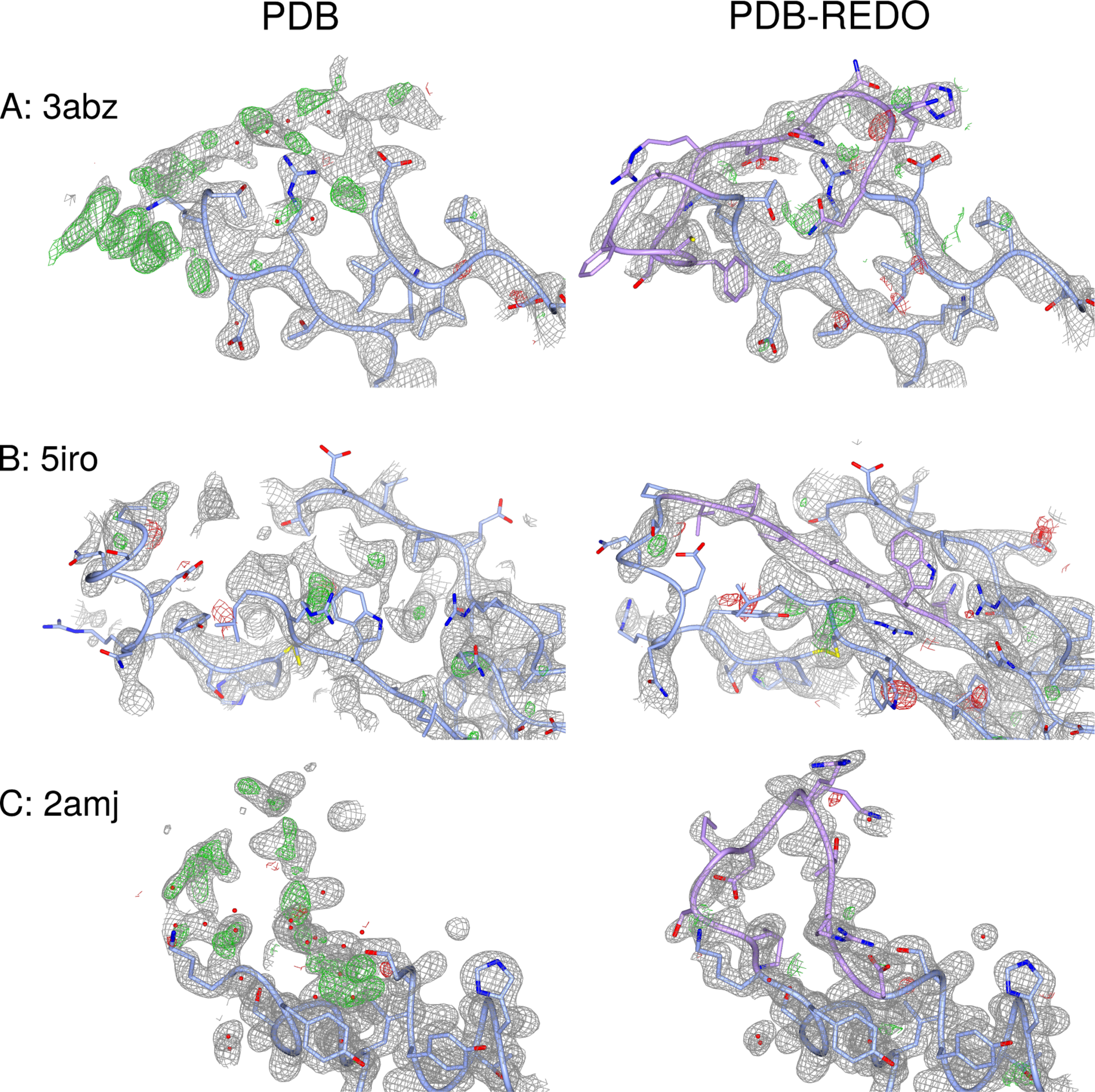
Examples of built loops. Newly built parts are shown in pink. (A) 3abz chain D residue 497-510. (B) 5iro chain I residue 243-249. (C) 2amj chain D residue 108-117. Details described in the main text. The 2mF_o_ - DF_c_ maps are shown at 0.8 σ, 1.2 σ and 1.0 σ, respectively. The mF_o_ - DF_c_ maps are shown at 3.0 σ in all cases.

A second example shows how the general improvement of fit to the crystallographic data in PDB-REDO can facilitate when loop building is included. As shown in Fig. 6B, a strand is missing from the β-sheet of the α3 subunit of HLA in PDB entry 5iro (Li *et al.*, 2016), a protein complex of Adenovirus type 4 E3-19K with HLA class I histocompatibility antigen. By optimization of the structure model by PDB-REDO including loops building, the R/R_free_ drops from 0.220/0.236 to 0.182/0.190. The overall improvement of the model leads to much clearer maps into which the missing strand can be built unequivocally.

Finally, two loops are missing from PDB entry 2amj, the apo-structure model of modulator of drug activity B (MdaB), a putative DT-diaphorase (Adams & Jia, 2006). In this paper, the authors discuss also the FAD-bound state structure (2b3d), where both loops are ordered: it is stated, that these loops become ordered upon binding of FAD due to a rearrangement in the hydrogen bonding network. However, there is clear density for one of the loops (L2). No less than 13 water molecules had been built into the density of the missing loop in chain D (Fig. 6C). It was proposed (Adams & Jia, 2006) that FAD binding induces loop stabilization through changes in the hydrogen bonding network; modeling this loop shows that the structure model of the apo-form and FAD-bound form are highly similar. Therefore the structural evidence does not necessarily support this claim.

## Discussion

The increasing number of residues that is not built in new protein structures can be attributed to many factors: the ever larger structures determined by X-ray crystallography (van Beusekom *et al.*, 2016) are more likely to contain flexible regions within stable scaffolds; better annotation of the sequence of the crystallized constructs (Henrick *et al.*, 2008; Dutta *et al.*, 2009) highlights missing regions better; and opportunities to built loops supported by the electron density are ignored due to haste or lack of experience (Mowbray *et al.*, 1999) as new generations of crystallographers are determining structure in a higher throughput. A worrying observation we made whilst teaching is that some students had deleted parts of a model to improve validation statistics (Read *et al.*, 2011) such as the percentage of RSRZ outliers.

We argue that models should be built as complete as possible given the data, because high completeness will increase their usefulness to the user community. For instance, simulations of protein complex formation might improve when the protein structure model is as complete (and correct) as possible: it is better to have a starting experimental conformation available even if supported only by weak electron density, rather than predicting it by purely computation methods. Also, the presence of loops in refinement improves the local structure quality of the loop surroundings because the added atoms impose better conformational restraints in the structural neighborhood.

In some instances, our methods were unable to build a missing loop in an area with relatively clear electron density. This had to do either with the lack of well-aligning homologous loops or with the poor geometry of the fitted fragments. In some cases, this could point to register errors (a few residues aligned erroneously onto the sequence). Better comparison between homologous structures, for instance using tools like *phenix.structure_comparison* (Moriarty *et al.,* 2017), could be used to identify such regions more reliably; this is not feasible in an automated fashion with current methods.

The algorithms we have developed to decide whether loops should be kept or not, may also be applied to existing parts of protein structure models. We have emphasized in this paper the amount of loops that is *not* built in the PDB but may be buildable; however, there may also be cases where crystallographers have modelled loops overenthusiastically. To estimate the extent of that, we analyzed the distribution of the density ratio between atoms in loops and random atoms in the template structures (the structures where the loops were copied from). This density ratio is better for the template loops than the density threshold cutoff for the newly built loops in most cases (Fig. 7): this is to be expected because the loops are normally missing precisely because their electron density was not very clear. However, there are also cases where the density fit of the template loop is quite poor: inspection of several cases shows that the majority of these loops have a likely correct conformation which is supported by the electron density; however, there are also cases where the loop should not have been modelled.

**Figure 7:**
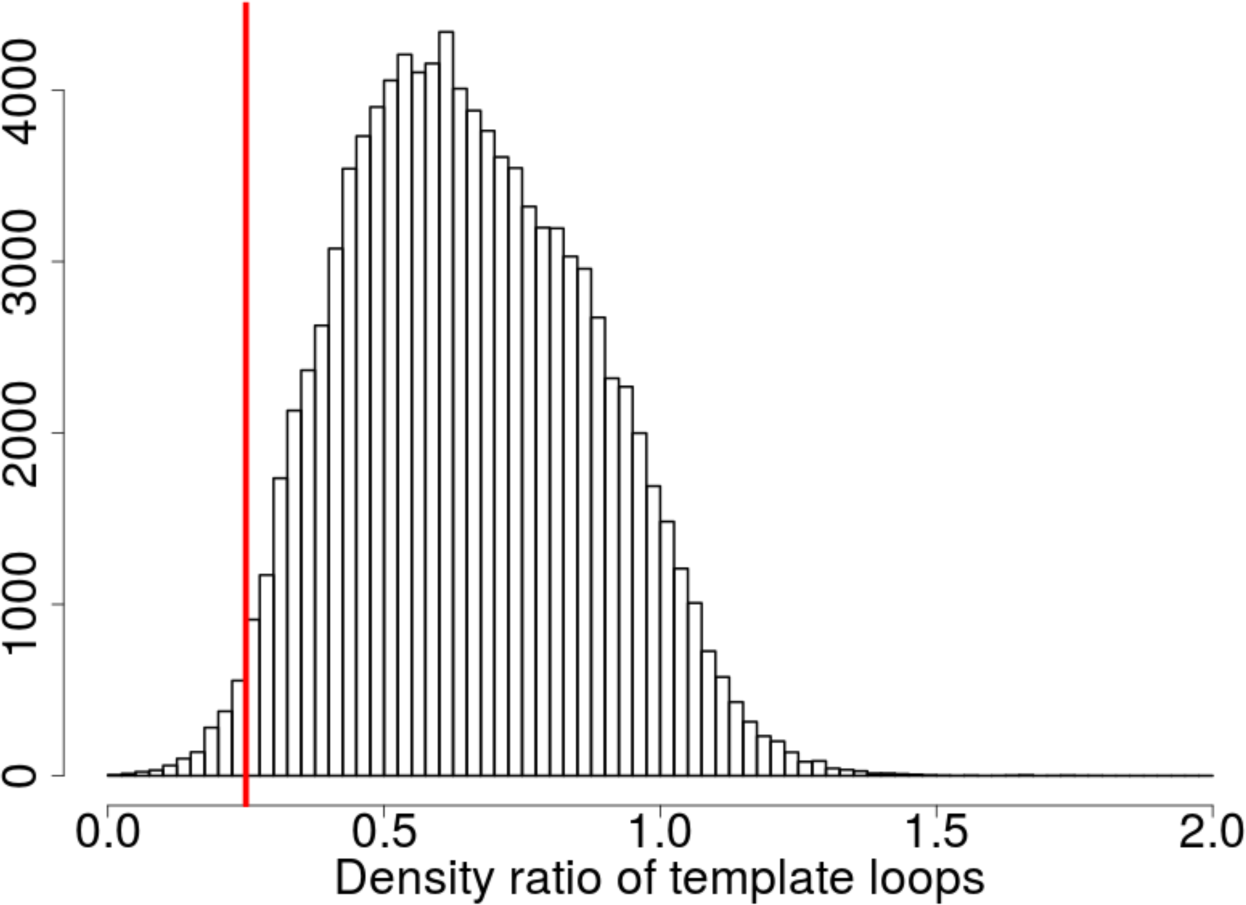
Frequency of specific density ratios of loop backbone versus the rest of the backbone for the loops that were used as template from homologous structures. Most template loops have sufficient density and therefore they would also have been built by *Loopwhole*: those loops are to the right side of the red line indicating the density ratio cutoff of 0.25.

A possible addition to the methods presented here is building partial loops or expanding the termini. At first, we allowed partial loops to be built by *Loopwhole* if the density fit for that part of the loop was above the density threshold. Although the majority of 8,217 partial loops we built was modelled correctly, building partial loops was not sufficiently reliable to automate. Too often, an amino acid would be modelled into neighbouring density of waters or unidentified ligands such as PEG. The same issue would arise for terminal extensions, with two additional difficulties. First, the number of residues that may be added is not always clear. Many residues can be missing from the terminus and, since it is *a priori* unclear how far the terminus can be extended, the residues should be added one by one. This is less efficient than loop fitting and therefore likely to cost much more CPU time per added residue. Second, instead of two anchor points on the ends of a missing loop, a terminus only has a single anchor point: the current terminus. The absence of a second anchor point means a drastic loss of information about the general direction in which the residues should be built. Therefore, it is more difficult to detect cases where the expanded fragment is not built into the correct electron density. The combination of these two limitations has kept us from implementing termini extension at present.

New structure models are added to the PDB every week, enriching the set of homologous structure data. The availability of suitable candidates for loop transfer will therefore only increase further, facilitating loop modeling for new structures. The pro-active updating of existing structure models by PDB-REDO ensures that older structure models also benefit from the increased availability of homologs. The original struggle to find a good loop conformation for the first published structure model in a protein family will remain, but it has become a temporary problem: solving the first structure of a protein provides a handle for much future structural research and now in return this future research also provides means to make this first structure more complete. Clearly, we have demonstrated that the increased availability of homologous data can be used to improve the completeness of protein structure models of the past, the present and the future.

## Availability

Both the PDB-REDO databank and web-server are hosted on https://pdb-redo.eu. On the web-server, crystallographers can submit their work-in-progress models to run PDB-REDO including the new loop building procedure. The 1,990 models from the test set are available through the databank. Existing databank entries are gradually updated to include the loop building procedure. On the PDB-REDO databank entry pages, registered users can submit an update request to prioritize the update of that entry. Binary executables of *Loopwhole, fixDMC*, and *tortoize* are available from the website and source code is available on request.

## Acknowledgements

This work is supported from the Netherlands Organization for Scientific Research (NWO) Vidi grant 723.013.003. Additional support was provided by Horizon 2020 programmes West-Life (e-Infrastructure Virtual Research Environment project No. 675858) and iNEXT (project No. 653706). The authors thank the NKI research high-performance computing facility for providing computational resources.

## Author Contributions

BvB designed and executed research, analysed all data, prepared figures and drafted the manuscript; KJ and MH contributed specific algorithms and code; RJ and AP designed and supervised the research.

